# *In situ* single-cell analysis of canonical breast cancer biomarkers: phenotypic heterogeneity and implications on response to HER2 targeting agents

**DOI:** 10.1101/2022.09.21.508826

**Authors:** Garazi Serna, Eloy García, Roberta Fasani, Xavier Guardia, Tomas Pascual, Laia Paré, Fiorella Ruiz-Pace, Antonio Llombart-Cussac, Javier Cortes, Aleix Prat, Paolo Nuciforo

## Abstract

Breast cancer is a heterogeneous disease. Tumor cells and the surrounding microenvironment form an ecosystem that determine disease progression and response to therapy. To characterize the breast cancer ecosystem and the changes induced by targeted treatment selective pressure, we analyzed 136 HER2-positive tumor samples for the expression of canonical BC tumor diagnostic proteins at a single cell level without disrupting the spatial context. The combined expression of HER2, ER, PR, and Ki67 in more than a million cells was evaluated using a tumor-centric panel combining the four biomarkers in a single tissue section by sequential immunohistochemistry to derive 16 tumor cell phenotypes. Spatial interactions between individual tumor cells and cytotoxic T cells were studied to determine the immune characteristics of the ecosystem and the impact on response to treatment. HER2-positive tumors displayed individuality in tumor cells and immune cells composition, including intrinsic phenotype dominance which only partially overlapped with molecular intrinsic subtyping determined by PAM50 analysis. This single cell analysis of canonical BC biomarkers deepens our understanding of the complex biology of HER2-positive BC and suggests that individual cell-based patient classification may facilitate identification of optimal responders or resistant individual to HER2-targeted therapies.

## INTRODUCTION

Breast cancer (BC) is a heterogeneous disease accompanied by differences in clinical, molecular, and biological features^1^, which creates a challenge for prognosis and treatment^2^. Currently, BC samples are stratified for clinical purposes based on tumor cells’ expression of ER, PR, HER2, and the proliferation marker Ki67. These immunohistochemistry (IHC) biomarkers together with clinicopathologic indexes are used to predict disease outcome^3^, for treatment decisions, and serve as surrogates for prognostic gene expression profiles (GEP)^4–7^ categorizing BC into four basic subtypes which are related – but not equivalent – to GEP-defined intrinsic subtypes^8^. Luminal A and luminal B are roughly equivalent to [ER+|PR+] HER2− and [ER+|PR+] HER2+ tumors, respectively, though a small percentage of [ER+|PR+] HER2− tumors with Ki67 positivity are reported to belong to the luminal B subtype^9^. HER2 enriched tumors refer to [ER−|PR−] HER2+ despite the different methods used on HER2 assessment. The [ER−|PR−] HER2− (also named triple negative tumors) subtype is mainly composed of basal-like tumors, which are highly heterogeneous including at least claudin-low^10^, metaplastic breast cancer^11^ and interferon-rich tumours^12^ in addition to core basal tumors as demonstrated by the accumulated evidence.

Although these stratifications have improved therapy success, patient responses vary within each subtype demanding better characterization of BC ecosystem. Targets of current therapies are heterogeneously expressed within and between patients. This heterogeneity equips cancer cells for proliferation, survival, and invasion and likely underlies differential treatment efficacies. This ecosystem is further shaped by cellular relationships (tumor cell-tumor cell, tumor cell-immune cell, …) and strategies targeting relationships that promote tumor development are promising.

Next generation technologies such as gene expression based molecular profiling and genetic testing are considered as the future of cancer diagnostics. Despite that, the results generated may be significantly affected by the level of intra-tumor heterogeneity in the bulk sample typically analyzed with those methodologies. Given the heterogeneity of cellular phenotypes and relationships, patients classification and treatment should ideally consider the entire tumor ecosystem. Recent single-cell RNA sequencing and mass cytometry studies provided hints into breast cancer complexity and how this may influence prognostic and response to treatment ^13–16^. However, no study specifically characterizes the distribution of common breast cancer biomarkers at a single cell level in HER2-positive breast cancer without disrupting the tissue architecture. In the present study, we explored the composition, heterogeneity, and spatial organization of HER2-positive breast cancer at a single-cell level resolution maintaining the spatial information as well as the treatment induced changes following dual HER2 inhibition with lapatinib and trastuzumab. To do so, we took advantage of an innovative technique recently developed in our lab which allows multiplex in situ biomarker analyses in a single FFPE tissue section. Customized algorithms were developed to extract the different phenotypic cell populations within the tissue for subsequent analyses. In addition, we analyzed the spatial distribution and interactions between tumor phenotypes, the interaction with immune cells and their impact in predicting response to treatment.

The results from this analysis might help understand better intra-tumor heterogeneity and improve treatment strategies.

## MATERIALS AND METHODS

### 1. STUDY POPULATION

Patients enrolled in the PAMELA phase II trial will be included in the study. Briefly, 151 patients with operable or locally advanced HER2-positive breast cancer were treated with neoadjuvant lapatinib (1000 mg daily) and trastuzumab (8 mg/kg IV loading dose followed by 6 mg/kg) for 18 weeks. Patients with hormone receptor (HR)-positive disease received letrozole or tamoxifen according to menopausal status. Formalin-fixed paraffin-embedded (FFPE) tumor samples at baseline and at D15 of treatment were collected according to protocol. Of the 151 patients enrolled in PAMELA study, 72 had a baseline sample and 64 had an on-treatment (day-15) sample for multiplexed immunohistochemistry analysis. Forty-nine patients had paired baseline and day-15 samples. Summary shown in **Supplementary Figure S1**.

From the all the patients, demographic patient data, tumor histopathological features (histotype, size, pT stage, pN stage), HER2 status (IHC and/or FISH results) and Hormone receptor status, PAM50 intrinsic subtype and treatment response data (pCR) were available. Clinicopathologic data is summarized in **Table 1**.

### 2. NGI (NEXT GENERATION IHC)

We used an automatized, simple, and flexible IHC protocol developed in our laboratory, named next generation immunohistochemistry (NGI) to study the expression of four canonical breast cancer biomarkers (ER, PR, KI67, HER2) at a single-cell level resolution^17,18^. The NGI protocol consists of iterative cycles of stanining/destaining on the same tissue section and uses the combination of Ventana Discovery Ultra (Roche Diagnostics), Nanozoomer slide scanner (Hamamatsu) and Visiopharm image analysis software. Briefly, an alcohol soluble chromogen (DISCOVERY AEC KIT (#760-258, Roche-Ventana)) was used to allow the destaining of the samples. After each automated IHC, samples were mounted in aqueous medium and digitalized (cycle 1). Subsequently, the section was destained in alcohol and submitted to the following staining cycle as shown in **Figure1A**. The sequence of the stainings were PR, KI67, HER2, ER followed by a PANCK staining for tumor area definition (antibody specification and protocol conditions in **Supplementary Table 1**.

**Figure 1.**
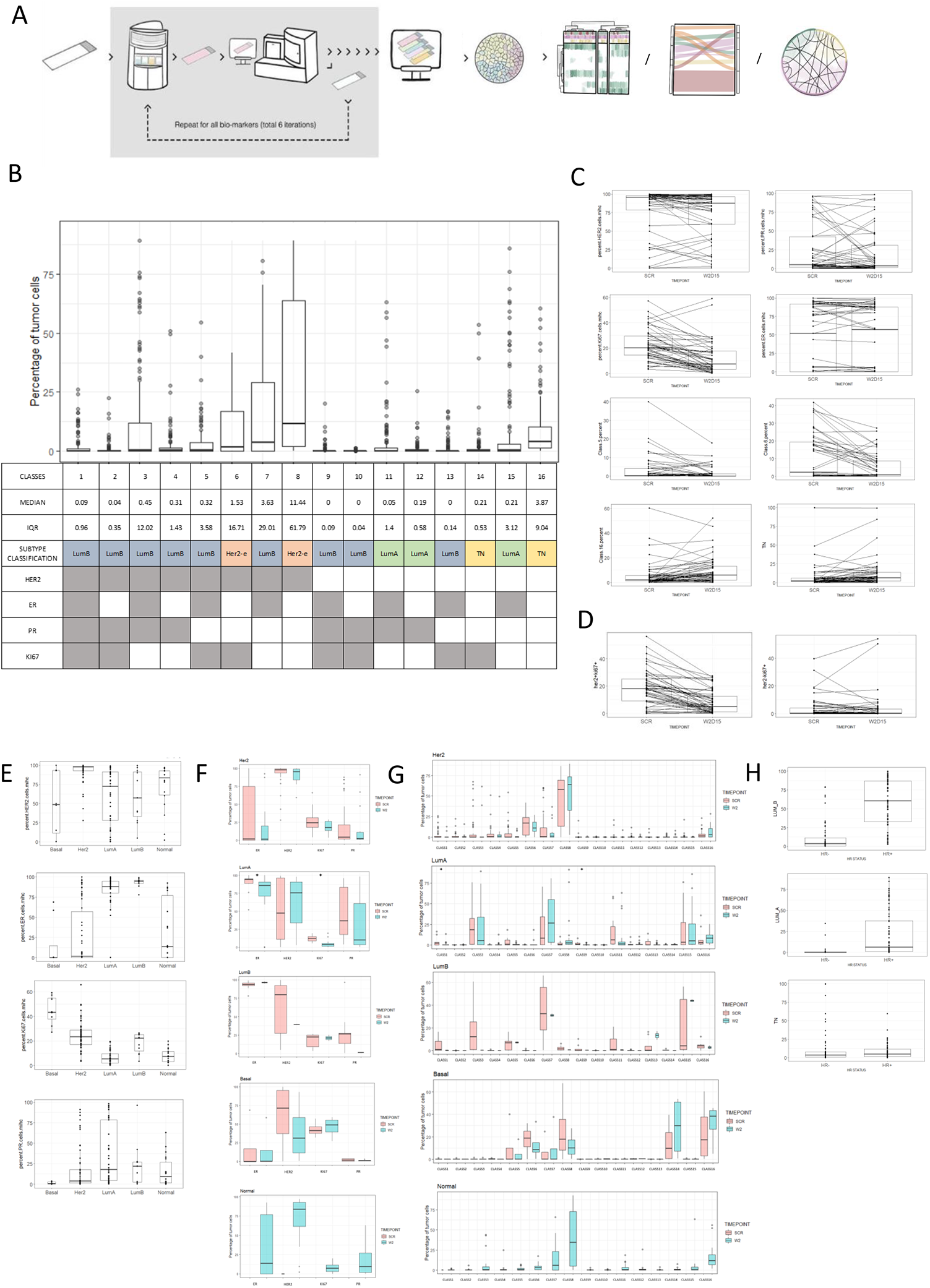
Her2-positive breast cancer composition. **A)** Illustration of NGI methodology used for the study. A single slide is stained for the first biomarker, scanned and destained and the process is repeated five times for each of the biomarkers. All obtained images are aligned and image analysis is used for the extraction of each tumor cell classification and location per sample. Information that is used for composition, clusterization and connectivity analyses. **B)** Box plot of the phenotype composition of her2-positive breast cancer samples, showing classes names, median, IQR for all samples and subtype classification into LumB, LumA, HER2-e and TN. In grey the positivity for each of the analyzed biomarkers (HER2, ER, PR and KI67) for each of the 16 generated classes. **C)** Box plots of phenotype-changes during the treatment in paired samples. **D)** Box plots of grouped Her2+KI67+ and Her2-Ki67+ changes during the treatment. **E)** Box plots of percentages of Her2, ER, PR and KI67 tumoral cells in the PAM50 groups. **F)** Box plots of general-composition of the different PAM50 groups. In pink SCR samples and in blue the day-15 samples. Star for significant differences between scr and day-15 samples medians. **G)** Box plots of phenotype-composition of the different PAM50 groups. In pink SCR samples and in blue the day-15 samples. Star for significant differences between scr and day-15 samples medians. **H)** Box plot of HR status differences in LumB, LumA and TN groups.

Before image analyses, individual images generated during each NGI staining cycle were aligned into a single virtual image using the Visiopharm® software. Once co-registered, image-analysis algorithms were applied to extract the data. First, a tissue recognition APP was run and then the PANCK staining was used to recognize the tumor for tumor analysis. A pathologist supervised the regions of interest. After that, a breast panel APP was run, which localizes and classifies all the tumor cells in positive or negative for ER, PR, KI67 and Her2 biomarkers. Any brown stain above the background level (average of 210 pixel-intensity) was considered positive for the classification. The number of each of the generated 16 tumor phenotypes (or classes) and the position of them is obtained by the APP. Image analysis algorithms are shared in supplementary material.

For subgroup analyses, the obtained classes were grouped into 4 categories (**Supplementary Table 2**): Her2-enriched (HER2E) for Her2-positive, hormone receptor negative cell phenotypes (classes 6 and 8), luminal A-like (LumA) for hormone receptor positive (ER -positive and/or PR - positive), Her2-negative, and Ki67-negative cells phenotypes (classes 11,12 and 15), luminal B-like (LumB) for hormone receptor positive (ER -positive and/or PR -positive) and HER2 - positive or KI67-positive cells (phenotypes 1-5,7,9,10 and 13), and triple negative (TN) for Her2-negative and hormone receptor negative phenotypes (classes 14 and 16).

For the neighborhood analyses, CD8 staining images generated on a consecutive section from our previous study^18^ were aligned with the PR images to extract tumor cell phenotypes and location. CD8 APP was run (image analysis algorithms are shared in **supplementary material**) and the number and location of each of them were obtained for consecutive analyses.

### 3. STATISTICAL ANALYSES

R software (v.3.6.1)^19^ was used for all statistical analyses. Statistical significance level was set to <0.05. Wilcoxon-Mann-Whitney non-parametric test was used for two group comparisons and Kruskal-Wallis test for three group comparisons. Heterogeneity analyses were done using vegan package. Intratumoral diversity (alpha diversity) was analyzed using richness, Shannon and evenness indexes. Intertumoral diversity (beta diversity) was analyzed using one of the most common metrics, the Bray-Curtis dissimilarity. Significance between groups was tested with Peranova test (adonis). Classes that had a median higher than 1% were used for the heterogeneity analyses. Hierarchical cluster analysis was done using pheatmap package and using Ward’s method to group patients with similar compositions. For neighborhood analyses, we connected the cells to each other by means of a Delaunay triangulation algorithm, using the centroid of the segmentation mask. This allowed us to locate the nearest neighbors avoiding those connections that are shielded by nearest cells. In addition, to avoid connections between cells that were too far apart, we established a maximum distance of 20 microns between two neighboring cells. The percentage of connections per sample was obtained and normalized by the total tumor cells for statistical analyses to ensure that tumor size was not affecting the results. The affinity of the CD8 to the different subtypes was calculated by dividing the percentage of the connections of the cd8 for each subtype by the percentage of that particular subtype in the sample.

## RESULTS

### 1. Single-cell composition analysis of HER2-positive breast cancer

We analyzed 136 prospectively collected tumor biopsies from the PAMELA trial, including 72 baseline and 64 day-15 samples from HER2-positive breast cancer using our NGI technology (**Supplementary Figure 1A**). Clinicopathological characteristics of the NGI cohort are summarized in **Table1**. In total, 1028974 cells were analyzed (mean=7566, median=4060, IQR=1378-9454). To ensure data quality, we compared NGI results generated with the matched IHC scores available as part of central confirmation analysis of the PAMELA trial. The frequencies of ER+, PR+, HER2+, and Ki67+ cells determined by NGI were comparable with the centrally determined pathological scores (**Supplementary Figure S2**).

**Table 1.**
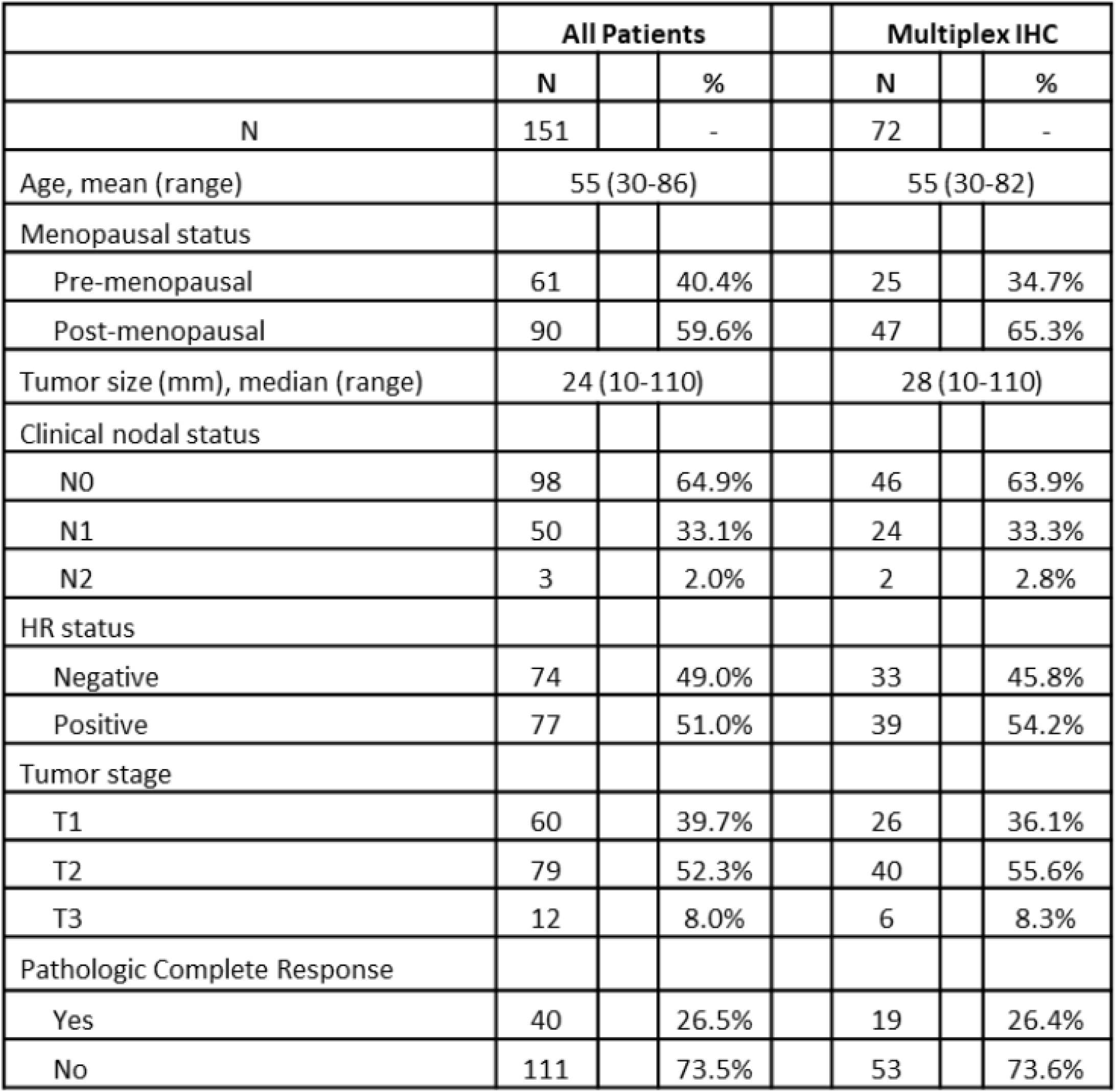
Summarized clinicopathologic data of the analyzed samples.

To characterize the composition of breast cancer at a single cell level, we used a tumor-centric panel including ER, PR, HER2, and Ki67 to classify individual tumor cells in 16 different phenotypes based on the combined markers expression of the 4 markers. (**Figure 1A, Supplementary Figure S2**). As expected, most cells in HER2-positive breast cancer were expressing HER2 (intrinsic cell phenotypes: 1-8, median=92.3%) followed by ER (intrinsic cell phenotypes 1,3,5,7,9,11,13 and 15, median=57.9%), Ki67 (intrinsic cell phenotypes 1,2,5,6,9,10,13 and 14, median=16.4) and PR (intrinsic cell phenotypes 1-4 and 9-12, median=5.7) (**Supplementary Table S3)**.

The analysis of distribution of individual intrinsic cell phenotypes revealed a heterogeneous distribution of tumor cell-intrinsic phenotypes in HER2-positive breast cancers, being the HER2E cell phenotype 8 the predominant cell phenotype (median=11.44, IQR=61.79) followed by TN phenotype 16 (median=3.87, IQR=9.04) and LumB phenotype 7 (median=3.63, IQR=29.01) (**Figure 1B, Supplementary Table S4**).

Analysis of paired samples showed an overall decrease of HER2-positive (median baseline=95.7%, median day-15=87.5%, p=0.022) and Ki67-positive (baseline=20.2%, day-15=7.5%, p<0.0001) cells from baseline to day-15. On the other hand, dual HER2 inhibition did not induce a significant shift in the overall composition of ER-positive and PR-positive cells. The overall decrease of HER2-positive and Ki67-positive cells from baseline to day-15 induced by the treatment was driven, at the intrinsic cell phenotype level, by the significant increase of non-proliferating TN cells (phenotype 16, median baseline=2.1%, median day-15=6.0%, p=0.0016) and decrease of proliferating HER2-positive cells (HER2+Ki67+, median baseline=18.1%, median day-15=5.1%, p<0.0001; phenotype 6, median baseline=2.4%, median day-15=0.9%, p=0.0001; phenotype 5, median baseline=0.3%, median day-15=0.01%, p=0.019) (**Figure 1C)**. Interestingly, no treatment effect was observed on proliferation in HER2-negative cells (median baseline=0.4%, median day-15=0.4%, p=0.6492) providing evidence that the treatment is specifically targeting Her2-positive cells (**Figure 1D)**.

The cell type frequencies varied among and between tumor molecular intrinsic subtypes determined by GEP, with a higher frequency of HER2-positive cells in HER2E samples (p-value<0.0001), PR-positive and ER-positive cells in luminal samples (p-value <0.0001), and proliferating KI67 positive cells in Basal samples (p-value <0.0001) (**Figure E, Supplementary Table S4)**.

At the individual intrinsic cell phenotype level, basal tumors showed a higher frequency of TN (phenotypes 14 and 16) and HER2E (phenotypes 6 and 8) cells compared to other breast cancer subtypes, which was statistically significant for TN phenotypes. Tumors of HER2E subtype were significantly enriched in HER2E cell phenotypes 6 and 8 with a lower frequency of the remaining intrinsic cell phenotypes compared to non-HER2E tumors. A higher frequency of luminal cell phenotypes was observed in luminal samples which were also showing significantly less HER2E cell phenotypes frequency than other tumor subtypes. LumB cell phenotype 5 significantly differentiates luminal A from luminal B tumors (median LumA=0.32, median LumB=6.77, p-value=0.0013). Normal samples were mainly composed of HER2E cells (phenotypes 6 and 8) and TN cells, the latter being significantly more abundant in normal compared to non-normal samples (median phenotype 16, normal=11.67, non-normal=3.22, p-value=0.00026) (**Supplementary Table S5)**. When comparing baseline with day-15 samples, we did not find significant differences neither at overall marker levels **(Figure 1F)** nor at individual intrinsic cell phenotype composition **(Figure 1G)** except for molecular intrinsic subtype luminal A where the percentage of ER+/Ki67+ and phenotypes 1 and 9 was significantly lower in on-treatment samples.

Lastly, we analyzed whether intrinsic cell phenotypes distribution varied according to the clinicopathological features of the tumor. As expected, clinical HR positive tumors showed a significantly higher proportion of ER-positive and PR-positive tumor cells as compared to HR-negative tumors and a lower proportion of KI67-positive and Her2-positive cells. Luminal intrinsic cell phenotypes were significantly enriched in clinical HR-positive compared to HR-negative tumors except for PR+/ER-phenotypes which were higher (phenotypes 2 and 4) or did not differ significantly (phenotype 10) in HR-negative tumors. HER2E intrinsic cell phenotypes (6 and 8) were also significantly enriched in HR-negative compared to HR-positive tumors (**Figure 1H, Supplementary Table S6**).

Tumor size and nodal stage did not significantly impact on the distribution of, neither the four biomarkers (**Supplementary Table S6**), nor the intrinsic cell phenotype tumor content.

### 2. HER2-positive breast cancer heterogeneity

Tumor heterogeneity is believed to drive disease progression or resistance to treatment. Intratumoral heterogeneity was determined using alfa-diversity indexes which quantify the number of different intrinsic cell phenotypes co-existing within a sample (richness index), their relative abundance (Shannon index), and how similar the phenotypes are numerically distributed (Pielou’s evenness index) within a sample. Intratumoral heterogeneity increased with tumor stage (Baseline eveness index: T1 vs T2, p=0.005; T1 vs T3, p=0.0027) indicating a progressive acquisition of different cell phenotypes with tumor growth. No significant differences were found according to nodal stage. Clinical HR-positive tumors were more heterogeneous than HR-negative tumors. This finding was in line with the higher heterogeneity observed in luminal tumors compared to other intrinsic molecular subtypes by GEP (**Figure 2A**). Treatment did not induce a significant shift in intratumoral heterogeneity as shown by paired samples analyses (median baseline=0.88, median on-treatment=0.84, p-value=0.52) (**Supplementary FigureS2A**). Intertumoral heterogeneity was determined using the Bray-Curtis matrix which quantifies the similarity of tumors based on intrinsic cell phenotypes composition and visualized with principal coordinates analysis plot (**Figure 2B**). Analysis of intertumoral heterogeneity failed to show significantly different compositions according to their tumor size or nodal involvement, with tumors segregating together independently of these clinicopathological features. Clinical HR-positive tumors segregated together and separately from HR-negative tumors (p-value <0.0001). This diversity was maintained during treatment in day-15 samples (p-value= 0.00316). Luminal tumors (A and B) clustered together and separately from Basal and HER2-E tumors. The difference was statistically significant at day-15 (p-value= 0.01769) with normal-like tumors clustering together with HER2E and Basal tumors. Intertumoral heterogeneity analyses on paired samples showed the same results (**Supplementary Figure S3)**.

**Figure 2.**
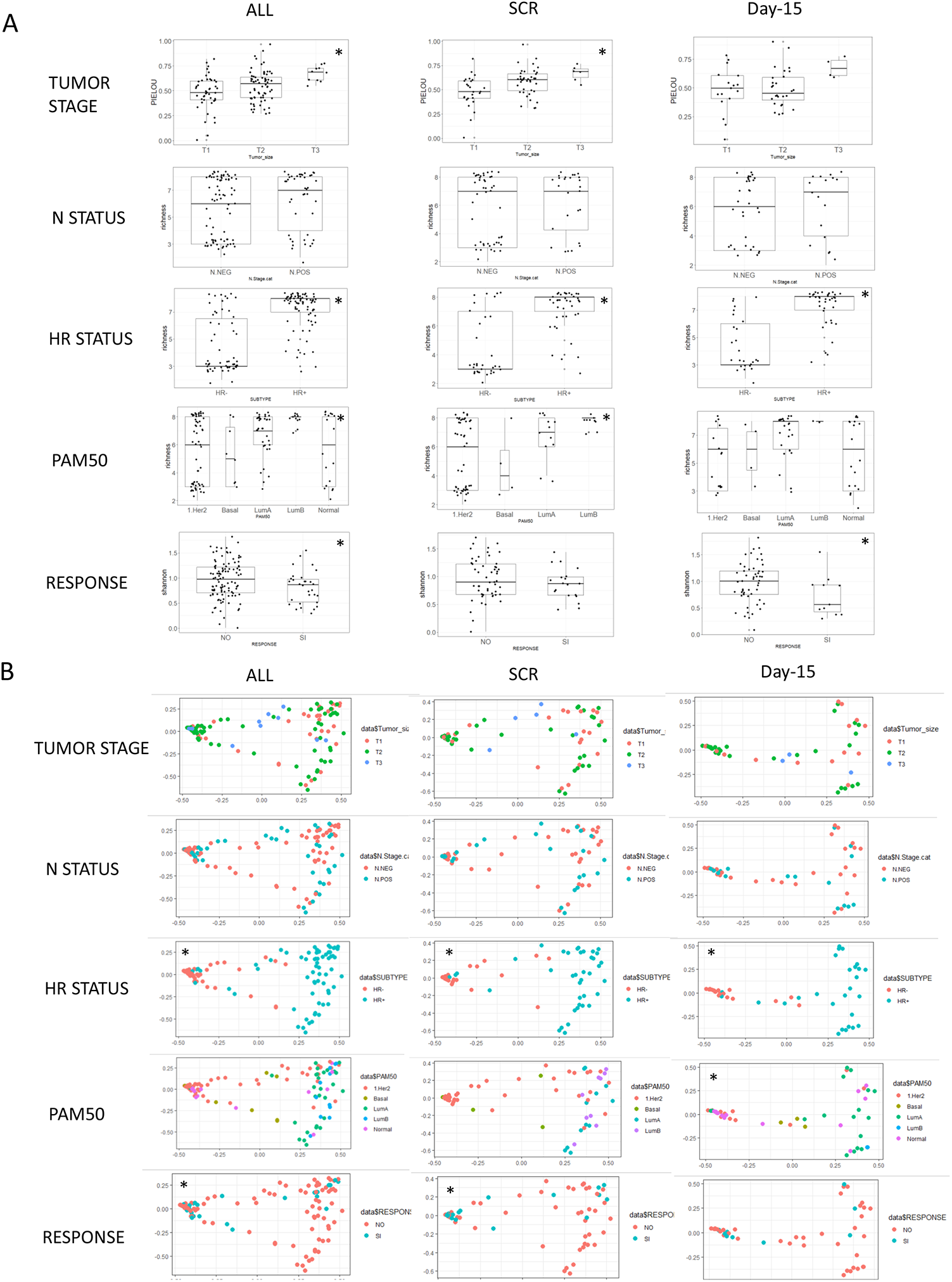
Heterogeneity analyses. A) Box plots showing intratumoral diversity variation for tumor stage, nodal status, hormone receptor status, pam50 classifications and response in all (left), scr (middle) and day-15 (right) samples. Star shows the significant intratumoral heterogeneity between groups. B) PCoA analysis plots of Bray-Curtis computed distances between all (left), scr (middle) and day-15 (right) samples. Different clinicopathologic features are shown in different colors. Star shows th e groups that are significantly different in phenotype composition.

Analysis of tumors according to response to anti-HER2 neoadjuvant therapy revealed that tumors from responders shared a similar composition at baseline (p-value=0.008) and separated from tumors from non-responders, suggesting that tumor intrinsic features predictive of pCR may be found before treatment is started. On the other hand, intratumor heterogeneity was not a tumor characteristic predictive of response at baseline but at day-15. Tumors from patients who did not achieve a pCR exhibited a significantly higher intratumoral heterogeneity at day-15 compared to tumors from patients who responded to the treatment (Shannon, p=0.044), indicating that on-treatment survival of different cell phenotypes may predict resistance (**Figure 2A**).

#### Response analysis by tumor intrinsic cell phenotype composition

To assess whether tumor intrinsic cell phenotype composition may affect response to neoadjuvant anti-HER2, we calculated whether the frequency of each individual tumor cell phenotype differed between patients achieving or not achieving pathological complete response (pCR).

As our dataset was a subset of the original PAMELA trial, we first determined whether this smaller subset was representative of the entire study in terms of response analyses. At baseline, TILS were significantly higher in responders (median=20) compared to non-responders (median =10, p=0.0017). At day-15, TILs (median responders=50, median non-responders=15, p=0.0036) and the CelTIL (median responders=58.3, median non responders=-1.5, p<0.0001) were significantly higher in responders thus confirming previous finding obtained from analysis of the full dataset.

Comparative analysis of tumor intrinsic cell phenotypes distribution between responders and non-responders did not show significant differences of any individual cell phenotypes between responders and non-responders both at baseline and day-15, except for LumB phenotype 2 at baseline (median responders=0.1%, median non-responders=0.03%, p=0.03), LumB class 11 at baseline (median responders=0.0%, median non-responders=0.04%, p=0.02), and LumB class 1 at day-15 (median responders=0.0% median non-responders=0.08%, p=0.04). However, median values for differentially abundant phenotypes were extremely low, being below 1% in both outcome groups **(Supplementary Figure S4**). When grouping the intrinsic cell phenotypes, we found a higher proportion of Her2-positive (median responders=95.9%, median non-responders=89.9%, p-value=0.028) and Her2-enriched (phenotypes 6 and 8) cells in responders (median=73.5%) as compared to non-responders (median=10.0%, p-value=0.047). Luminal A phenotypes (11,12 and 15) were enriched in non-responders (median non-responders=1.3%) as compared to responders (median=0.2%p-value=0.003) **(Figure 3A**).

**Figure 3.**
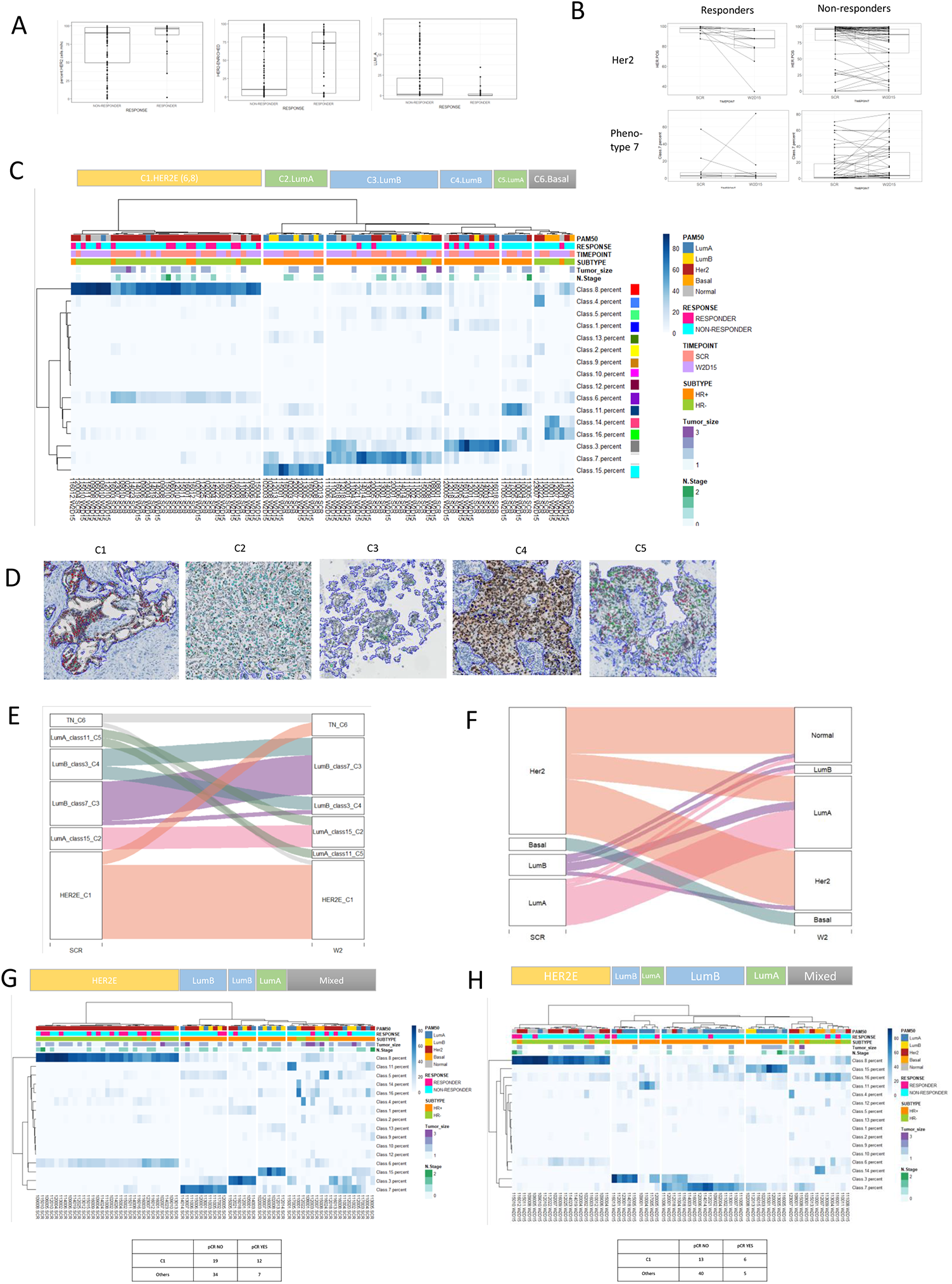
Response analyses. **A)** Boxplots of significant variables between responders and responders. **B)** Boxplots of treatment induced changes in paired responders and non-responder patients. **C)** Heatmap showing the percentage of each phenotypes of paired scr and day-15 samples. Each cluster is generated with patients with similar phenotype compositions. Heatmap colors represent percentage of each of the phenotypes. Annotation colors are shown next to the heatmap. **D)** Representative images of classified tumors from each cluster. Colors of the cells are shown next to the classes in the heatmap. **E)** Alluvial plot showing the cluster changes of paired patients during the treatment. **F)** Alluvial plot showing the Pam50 group changes during the treatment. **G)** Heatmap of SCR samples. Number of responders and non-responders are shown for the first cluster and for the others. **H)** Heatmap of day-15 samples. Number of responders and non-responders are shown for the first cluster and for the others.

The total number of HER2-positive cells significantly decreased in patients responding to anti-HER2 (median baseline=97.9%, median day-15=87.6%, p=0.009) but not in patients who did not, suggesting a reduction in tumor burden. On the other hand, patients who did not achieve a pCR showed an increase of LumB phenotype 7 with treatment (median baseline=0.8%, median day-15=3.9%, p=0.042) which was not observed in responders, possibly reflecting the expansion of a resistant clone (**Figure 3B**).

Consensus clustering was performed to group samples from HER2+ patients according to the distinct cell phenotypes. Unsupervised hierarchical clustering (**Figure 3C**) using paired samples classified tumors into six groups (C1-C6). Cluster 1 (HER2E) was dominated by tumor cells from phenotypes 6 and 8. Tumors in cluster 1 were clinical HR-negative (86.1%) and molecular intrinsic subtype Her2-E or Normal-like (94.7%). Clusters 2 and 5 (LumA) were enriched in cells from phenotypes 15 and 11, respectively. All tumors were clinical HR-positive. 63.2% were LumA and 21.1% LumB by intrinsic molecular subtyping. Clusters 3 and 4 (LumB) were composed mostly of cells from phenotypes 7 (cluster 3) and 3 (cluster 4). All but two tumor samples in these clusters were clinical HR-positive and exhibited a mixed molecular intrinsic subtyping (32.3% Her2-E, 41.2% Lum-A, 8.8% Lum-B,5.9% Basal and 11.8% Normal). Lastly, cluster 6 (Basal) was enriched in TN phenotypes 14 and 16. Tumors were clinical HR-negative (7/8, 87.5%) and half exhibited a basal intrinsic molecular subtype. Baseline and on-treatment samples from the same patients tended to group together in the same cluster or moved to a similar cluster during treatment (**Figure 3E**) More treatment-induced changes were observed in molecular intrinsic subtyping by PAM50. Similarly, only 20% of LumB tumors remained LumB after treatment. Treatment did not significantly impact on LumA and Basal tumors which tended to maintain the same subtype after treatment (**Figure 3F**).

To determine whether patients responding or not responding to anti-Her2 therapy segregated within a specific cluster, we performed consensus clustering of all (paired and unpaired) baseline and on-treatment samples independently. At baseline, hierarchical clustering classified patients into five groups corresponding to HER2E (C1, phenotypes 6 and 8), LumB (C2 and C3, phenotypes 7 and 3), LumA (C4, phenotype 15) and mixed Luminal/Basal (C5) clusters. Twelve out of 31 (39%), 3 out of 16 (19%), 0 out of 15 (0%) and 4 out of 19 (20%) patients from HER2E, LumB, LumA, and mixed clusters achieved a pCR, respectively. Patients in the HER2E cluster C1 had a significantly higher probability of responding to anti-HER2 therapies as compared to those in other clusters (Fischer exact test, p=0.01) (**Figure 3G**). At day-15, clustering analysis showed six tumor clusters with similar phenotype compositions as the baseline clusters. Six of 19 patients (32%), 2 out of 21 (9.5%), 1 out of 12 (8%), and 2 out of 12 (17%) patients from HER2E, LumB, LumA, and mixed clusters achieved a pathological complete response, respectively. Patients in the HER2E cluster had a non-significant higher probability of responding to anti-HER2 therapies as compared to those in other clusters (Fischer exact test, p=0.06) (**Figure 3H**).

### 3. Breast cancer tumor and immune cells relationship

Interactions between tumor cells and immune cells within the tumor microenvironment drive disease progression and response to treatment. To study homotypic and heterotypic relationships between tumor and immune cells, we determined the percentage of connections between each individual tumor cell phenotype, the distribution of tumor cell phenotypes according to tumor infiltrating lymphocytes (TILs) and the spatial relationship between cytotoxic (CD8+) immune cells with each tumor cell phenotype.

Homotypic tumor cells relationships were the most common. Most of the connections were found between cells belonging to the same or similar phenotype. Most of the homotypic connections were found between Her2E phenotype 8 cells, followed by connections between LumB phenotype 3 cells, TN phenotype 16 cells, LumA phenotype 11 cells and heterotypic connections between phenotype 8 with phenotypes 6 and 7 (**Figure 4A**).

**Figure 4.**
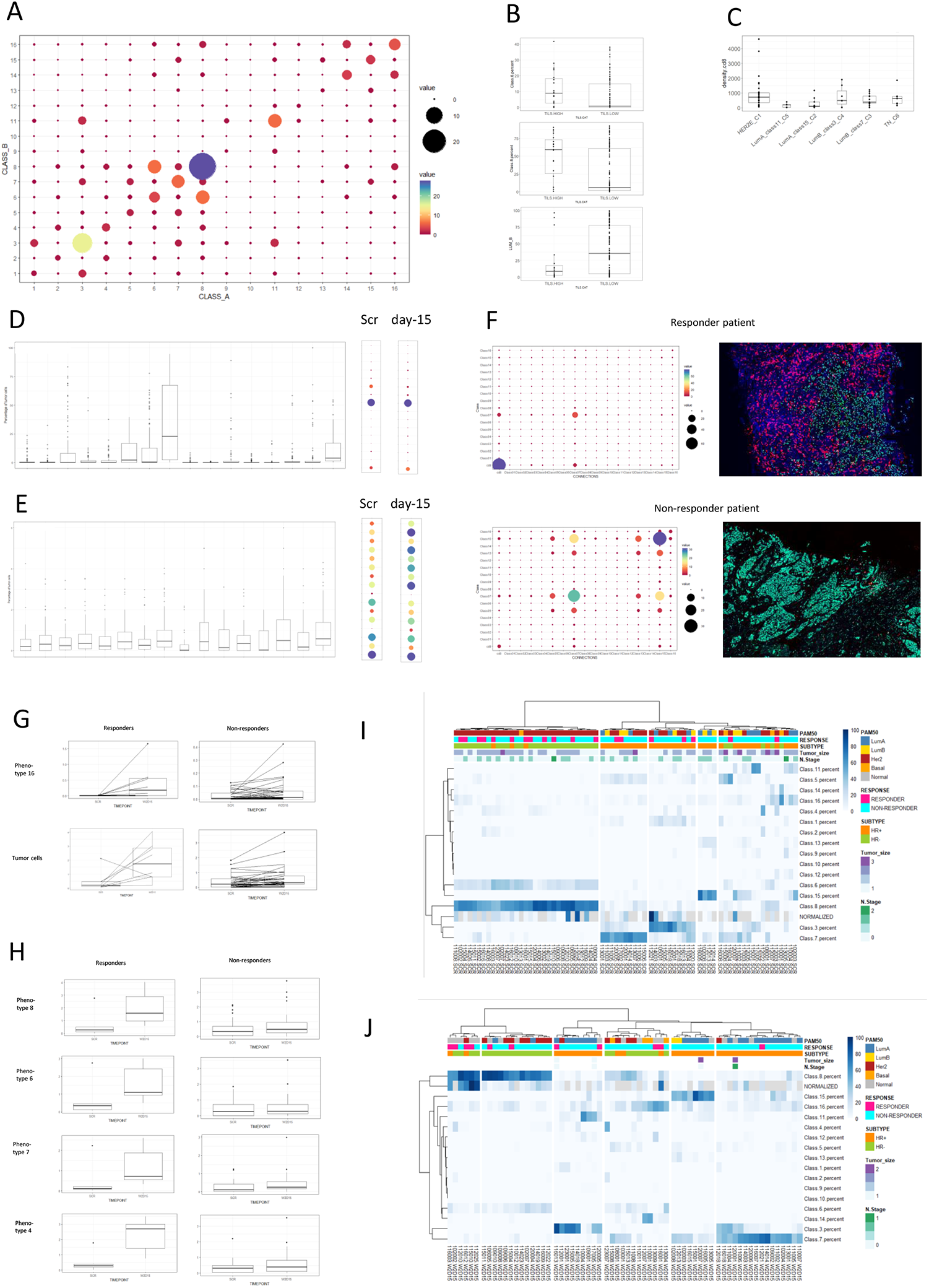
Spatial analyses. **A)** Dot-plot showing the connections between different tumor phenotypes. The size and color of the dots represent the proportion of connections. **B)** Boxplot of significantly different variables between TIL high and TIL low groups. **C)** Box plot of CD8 densities in the different clusterized groups’ samples. **D)** Box plots of the CD8 connections with each of the tumor phenotypes in all samples. Dot plots of the medians in SCR and day-15 are shown in the right. **E)** Box plots of the CD8 affinity with each of the tumor phenotypes in all samples. Dot plots of the medians in SCR and day-15 are shown in the right. **F)** Representative dot plots and virtual multiplexed images of a responder and non-responder patient. The size and color of the dots represent the proportion of connections between CD8 and tumor cells. CD8 cells are shown in red in the image. **G)** Boxplots of treatment induced connection-changes in paired responders and non-responder patients. **H)** Boxplots of treatment induced affinity-changes in paired responders and non-responder patients. **I)** Heatmap of SCR samples with CD8-tumor connection information. Annotation colors are shown next to the heatmap. **J)** Heatmap of day-15 samples with CD8-tumor connection information. Annotation colors are shown next to the heatmap.

To determine whether tumor cell phenotypes were differentially enriched in breast cancer according to the level of immune infiltration, we correlated the proportion of TILs with that of each individual cell phenotype in the same sample. Levels of TILs were positively correlated with HER2E phenotypes 6 and 8 both at baseline (0.49 and 0.41, respectively) and at day-15 (0.32 and 0.55, respectively) (**Supplementary Figure S5 A**). When we grouped tumors into inflamed (TILs >50%) and non-inflamed (TILs <50%) based on TILs infiltration, inflamed tumors were again significantly enriched in HER2E cell phenotypes 6 (median high=8.9%, median low=0.7%, p=0.026) and 8 (median high=59.3%, median low=6.3%, p=0.005) and depleted of LUM-B cells (median high=8.9%, median low=35.6%, p=0.048) compared to non-inflamed tumors (**Figure 4B, Supplementary Table S7**).

After finding the enriched subtypes in inflamed tumors, we wanted to explore whether TILs levels differed according to tumor intrinsic cell phenotype-defined clusters. We found that patients from clusters 2 and 5 (non-responder clusters) had significantly lower cd8 densities (median cd8 density in cluster 2= 114.5cells/mm2 and cluster 3 202.2 cells/mm2) while cluster 1 patients (responders) had the highest cd8 densities (median 723.5 cells/mm2) **(Figure 4C)**.

To determine the spatial relationship between CD8+ immune cells and tumor cell phenotypes, we calculated the proportion of intratumoral immune-tumor cells connections. The most common connections were between CD8 and HER2E tumor cell (phenotypes 8, median=22.75%; phenotype 6, median=2.5%) followed by TN cell phenotype 16 (median=3.93%). Similar results were found when dividing the dataset into baseline and day-15 samples **(Figure 4D, Supplementary Figure S5 B)**. Anti-HER2 treatment induced a general increase of number of connections between CD8+ immune cells and tumor cells (median baseline=0.2%, median day15=0.5%, p=0.01) without significant changes in connections of CD8 with any individual tumor cell phenotypes **(Figure 4D, Supplementary Table S8)**.

The higher number of connections observed between CD8 immune cells and HER2E and TN tumor cells phenotypes may be attributable to the higher prevalence of these phenotypes within the tumor. To elucidate whether cytotoxic T cells tended to interact preferentially with a specific tumor cell phenotype within a tumor, we determined the affinity of cytotoxic T cells for each tumor cell phenotype by dividing the percentage of connections of CD8+ cells with each of the phenotype by the percentage of tumor cells that were of the same phenotype.

More affinity of CD8 was found with TN cell phenotype 16 (median=0.58) followed by TN cell phenotype 14 (median=0.50), Her2E phenotype 8 (median=0.47), LumB phenotype 10 (median=0.44) and Her2E phenotype 6 (median=0.43) **(Figure 4E)**. The affinity of CD8 for the Her2E cell phenotype 8 (median baseline=0.3%, median on-treatment=0.6%, p=0.018) and LumB phenotype 7 (median baseline=0.1%, median on-treatment=0.3%, p=0.029) significantly increased with treatment (**Supplementary Table S9**).

Lastly, we wanted to analyze whether differences in the types of connection and affinity between responders and non-responders could be found. Overall, no significant differences in the number of connections between cd8 and the different cell phenotypes were observed between responders and non-responders. At baseline, no significant differences of CD8 connections or affinity were found between responders and non-responders. On day-15, tumors from patients responding to the treatment showed an overall increase in CD8/tumor cell ratio, connections of CD8 with any tumor cells, with HER2-positive cells and with phenotype 8 cells compared with tumors from non-responders. Similarly, higher CD8 affinity for HER2E phenotypes (phenotype 6 and 8), LumB phenotypes (4, 7, and 13) and LumA phenotype 11 was found in responders compared to non-responders (**Figure 4F, Supplementary Table S10**). These findings were confirmed in paired sample analysis.

Anti-HER2 treatment induced a significant increase in the number of CD8 connections with any tumor cell (median baseline=0.2%, median on-treatment=1.7%, p=0.007) and, specifically, with TN phenotype 16 (median baseline=0.0%, median on-treatment=0.2%, p=0.02) in responders compared to non-responders **(Figure 4H)**. Similarly, affinity for the Her2E class8 (median baseline=0.3, median on-treatment=1.6, p=0.0059) and class6 cells (median baseline=0.4, median on-treatment=1.5, p=0.022); LumB class7 (median baseline=0.1, median on-treatment=0.7, p=0.04) and class4 cells (median baseline=0.3, median on-treatment=2.7, p=0.013) **(Figure 4I)** increase in the responders but not in non-responders.

Due to the higher number of connections between CD8 and tumor cells observed in responders compared to non-responders, we included this feature to the intrinsic tumor cell phenotypes and performed unsupervised clustering analysis. The addition of immune feature to tumor features did not improve segregation of responder vs non-responders at baseline (**Figure 4J**). However, at day-15, the number of connections between cytotoxic T cells and tumor cells separated the HER2E cluster in two distinct clusters with different response rate the one with more connections specially enriched in (67% of responders) as expected as compared to the other her2 enriched cluster that became not enriched in responders (15% of responders) (**Figure 4K**).

## DISCUSSION

In this study, we conducted an *in situ* single-cell analysis of Her2 positive breast cancer using a sequential immunohistochemistry protocol combined with image analysis to provide virtual multiplexed expression of different biomarkers on the same slide. The developed methodology (called NGI or next generation immunohistochemistry) provided information on the expression of 4 canonical breast cancer biomarker at an unprecedented resolution allowing analysis of their co-expression in the same cell, their distribution, their spatial interaction (tumor cell-tumor cell, tumor cell-immune cells) and their changes during anti-Her2 treatment.

ER, PR, Her2 and Ki67 are standard biomarkers used in clinical practice for diagnosis and prognostication of breast cancer. Their expression is determined by conventional immunohistochemistry and their status defined by consensus cutoffs recommended in clinical guidelines. HER2-positive tumors are defined by >10% tumor cells showing strong (3+) complete membrane positivity or equivocal 2+ weak to moderate complete membrane staining in >10% of the tumor cells with Fish-negative result^20^. Similarly, hormone receptor status is defined by >10% tumor cells showing positivity at any intensity of ER and/or PR. For Ki67, no standard cut-off exists. A 20% threshold is generally used to define a high proliferative tumor and usually associated with poor prognosis^21–23^.

Treatment strategies are currently defined according to the individual status of each one of these biomarkers. However, treatment responses are heterogeneous and a better understanding of the breast cancer tumor ecosystem is needed.

Transcriptomics and proteomics approaches have been used to analyze the landscape of breast cancer and dig into the complexity of the tumor. However, these studies are performed on bulk samples, thus lacking contexture analyses^24–26^. Single-cell studies have provided a deeper understanding of the different cell populations distribution, tumor heterogeneity and influence on cancer progression and resistance^27–32^, but to the best of our knowledge, no studies have performed single cell level resolution analyses in Her2-positive breast cancers during the anti-her2 treatment.

The approach we described in this study combines a routine, widely used methodology such as IHC with image analyses algorithms to extract complex data from a single slide. Despite immunohistochemistry is traditionally considered a qualitative, single-plex, low-throughput methodology, our NGI protocol allows to virtually multiplex up to 6 (or more) different antibodies on the same slide to simultaneously study the co-expression, distribution, spatial interactions, and function of biomarkers of interest in the target cell (tumor or immune) with a single-cell level resolution and without disrupting the tissue organization by maintaining the spatial information. In this study, we developed a custom-made 5-plex panel which included HER2, ER, PR, Ki67 (classical breast cancer biomarkers) and CK (as tumor mask) to classify more than million cells into 16 different tumor cell intrinsic phenotypes. With an average of 4060 cells/sample analyzed; our approach is at least equivalent (if not superior in term of cells analyzed) to other single-cell breast cancer studies^30–34^.

Cell phenotypes prevalence and distribution were investigated in HER2-positive breast cancer from the PAMELA trial and correlated with clinicopathological and molecular features of the tumors. Intratumoral and intertumoral heterogeneity was quantified and impact of anti-HER2 inhibition on the cellular composition of the tumors and response to treatment investigated. Finally, tumor cell phenotype information was integrated with cytotoxic T cells spatial data to study homotypic and heterotypic relationships between tumor and immune cells and how interactions within the tumor microenvironment may predict treatment response or resistance.

Several confirmatory findings validated our experimental approach. First, comparative analysis of biomarkers expression determined by NGI was highly correlated with those obtained by standard immunohistochemistry analysis performed at central laboratory. Second, the overall reduction of Her2 and ki67 from baseline to day-15 confirmed previous findings reported in the Pamela trial using alternative methodologies as well as those from other neoadjuvant studies in Her2 positive breast cancer. Importantly, higher resolution analysis of tumor individual cell populations showed that Her2-positive breast ecosystem is constituted by multiple tumor cell phenotypes (both her2+ and her2-), which are differently modulated by the treatment. Non proliferating Her2+/ER-/PR-represents the predominant phenotype (class 8) followed by triple negative (16) and luminal B (7) phenotypes. Dual her2 blockade did not affect all phenotypes in the same manner. The significant reduction of proliferative her2 positive phenotypes-only without an effect on her2 negative ones is indicative of a clear pharmacodynamic effect of dual her2 inhibition^24,25,35^.

Poor prognosis and therapy resistance are associated with tumor heterogeneity^36,37^. We analyzed intratumor and intertumor heterogeneity according to breast cancer intrinsic cell phenotype composition and their impact on response to treatment. First, we found that intratumoral heterogeneity increases with tumor stage indicating a progressive acquisition of different cell phenotypes with tumor growth^38^. Second, we found that HR-positive tumors (clinical and luminal subtype by PAM50) exhibited higher intratumor and intertumor heterogeneity as compared with HR-negative tumors (clinical HR-negative and HER2E by PAM50). Third, tumor heterogeneity was inversely correlated with the probability of achieving a pCR. These findings may explain why her2-enriched and HR-negative patients, which are homogeneously composed predominantly by tumor intrinsic cell phenotypes 8 and 6, showed the highest rate of response to anti-HER2 treatment^24,25^.

On the other hand, clinical HR-positive, luminal tumors by PAM50 were highly heterogenous in composition with significant enrichment of different LumB (3 and 7) and LumA (11, 12 and 15) intrinsic cell phenotypes and depletion of HER2E phenotypes (6 and 8) which, in turn, resulted in reduced sensitivity to HER2 targeted therapy. Interestingly, the association found between higher intratumoral heterogeneity at day-15 and lower response rates indicated that on-treatment survival of different cell phenotypes may predict resistance. Heterogeneity analysis also revealed that tumor intrinsic features predictive of pCR may be found before the treatment is started as shown by the similar ecosystem found at baseline in tumors from responder with beta diversity analysis.

Consensus clustering was performed to group samples from HER2-positive patients according to the distinct cell phenotypes composition and correlated with response. Six different clusters (one HER2E, 2 LumA-like, 2 LumB-like, and one Basal) were identified based on tumor intrinsic cell phenotypes composition which only partially recapitulated molecular intrinsic subtyping by PAM50. Baseline and on-treatment samples from the same patients tended to group together in the same cluster or moved to a similar cluster during treatment. In contrast, PAM50 changes during treatment were notable with 20% of HER2E tumors shifting to LumA molecular subtype. Patients in the HER2E cluster (phenotypes 8 and 6) had a higher probability of responding to anti-HER2 therapies as compared to those in other clusters (Baseline, p=0.01; day-15, p=0.06). Importantly, patients in LumA clusters (enriched in phenotypes 11 and 15) were exquisitely resistant to anti-HER2 inhibition. This data provides more insight into our previous observation that HER2E tumors cells that are sensitive to anti-HER2 therapy but do not die acquire a Luminal A phenotype^35^. Our new results point to tumor intrinsic cell phenotype 15 (ER+/PR-/HER2-/Ki67-) as a resistant clone which pre-exist at low frequencies in HER2-positive breast cancer even before treatment starts, independently of the PAM50 subtype. The observed rapid acquisition (after 14 days of treatment) of PAM50 LumA molecular subtype may indeed reflect the expansion (as consequence of reduction of HER2E cells due to treatment) of this preexisting resistant clone. On the other hand, PAM50 HER2E tumors that became normal-like, an on-treatment biomarker of tumor responsiveness, are still predominantly composed of HER2E cells (phenotypes 8 and 6) which are diluted by stromal contamination due to decrease in tumor burden as consequence of response.

We also found LumB phenotype 7 (HER2+/ER+/PR-/Ki67-) as a resistant phenotype. This finding reflects the possible effect of the strong interplay between HER2 and ER, which may negatively impact on the response to anti-HER2 ^27,30,39,40^.

The study of how tumor and immune cells interact within the tumor microenvironment may increase our understanding of treatment response^25,41^. We’ve previously reported that patients responding to dual anti-HER2 inhibition showed higher TILS compared to non-responders^18^. In the present study, we further dig into tumor-immune cells relationship by spatial analysis. Inflamed tumors were enriched in HER2E cells (phenotypes 6 and 8) as opposed to non-inflamed ones where Luminal phenotypes predominated. In line with these observations, densities of CD8+ immune cells were higher in the HER2E cluster with LumA cluster showing the lowest levels of cytotoxic T cell infiltration. Connectivity analysis showed a higher number of connections between CD8+ immune cells and HER2E and TN tumor cells phenotypes as compared to other phenotypes, more affinity of CD8 for TN cells (phenotypes 14 and 16) and less affinity for hormone receptors positive cells in general. This observation matched with the differences in immunogenicity described according to breast cancer subtypes and partially explains why HER2-positive and TN tumors are more immunogenic^42^ as well as why HER2-positive breast cancer enriched in HER2E responded significantly better to anti-HER2 inhibition compared to other phenotypes.

During her2-treatment, cytotoxic T cells increase significantly^18,24,25^ and higher densities of intratumoral CD8s associated with pCR^18^. Here, we now showed that this increase reflects a higher number of connections and increased affinity of CD8 with tumor cells and HER2-positive cells in responders which is not found in patients who did not respond to the treatment. Lastly, the addition of the immune microenvironment status to tumor intrinsic cell phenotypes separated the HER2E cluster at day-15 into two subgroups with different response rates based on the connections between CD8 and tumor cells.

One of the biggest limitations of the study is the limited number of cases which did not allowed us to develop a predictive model of response to the treatment. Second, the number of biomarkers analyzed simultaneously on the same section is lower as compared with other single-cell, high throughput methodologies. Third, our study did not consider the different levels of expression of HER2, which is particularly relevant now due to the development of antibody-drug conjugates targeting the HER2-low breast cancer population. Future studies would be needed to address these gaps.

In conclusion, we have analyzed the HER2-positive breast cancer ecosystem at a single-cell resolution using a relatively simple and inexpensive methodology developed in our laboratory. Our findings confirmed previous observation using more expensive and less accessible approaches and expand our knowledge of Her2-positive breast cancer biology and composition during anti-her2 treatment providing potential clinically important information that could be used to improve targeted therapies.

## ACKNOWLEDGEMENTS

We thank all the patients and family members for participating in the PAMELA study. Acknowledgements to the Cellex Foundation for providing research facilities and equipment. Acknowledgements to Roche Diagnostics and Ventana Medical Systems, Inc. for providing the beta DISCOVERY AEC KIT reagent for research purposes. This work has been realized in the Surgery and Morphological Sciences Doctorate framework of Univ Autonoma de Barcelona. Eloy García holds a Juan de la Cierva-training grant (ref.FJC2019-040039-I).

## FIGURE LEGENDS

**Supplementary Figure 1.**
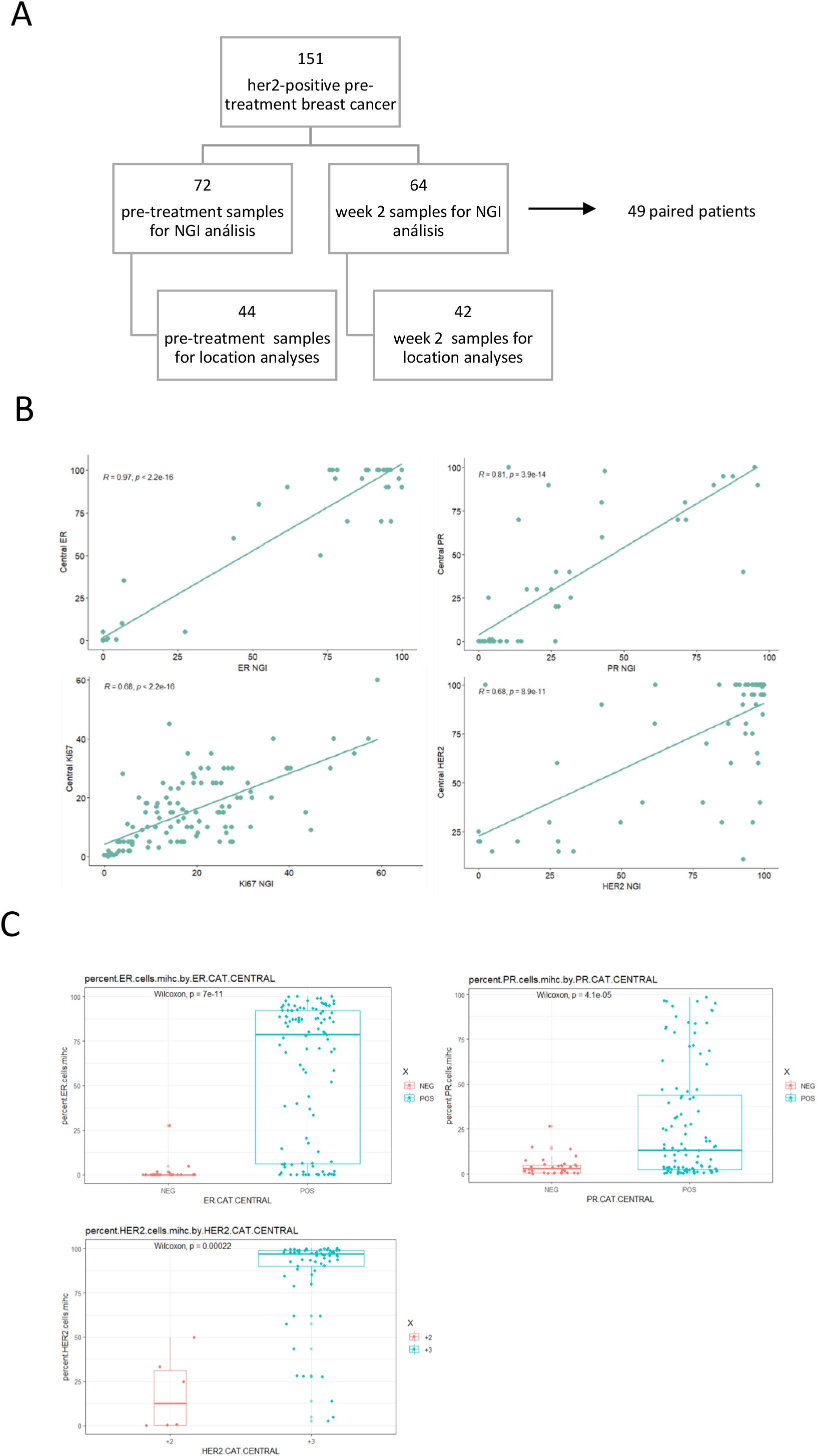
**A)** Sample cohort diagram. **B)** Scatter plot and Spearman coefficient of NGI and central lab results for each of the 4 canonical biomarkers. **C)** Box plots of the percentage of positive cells for the different biomarkers with NGI for the central lab categorized results.

**Supplementary Figure 2.**
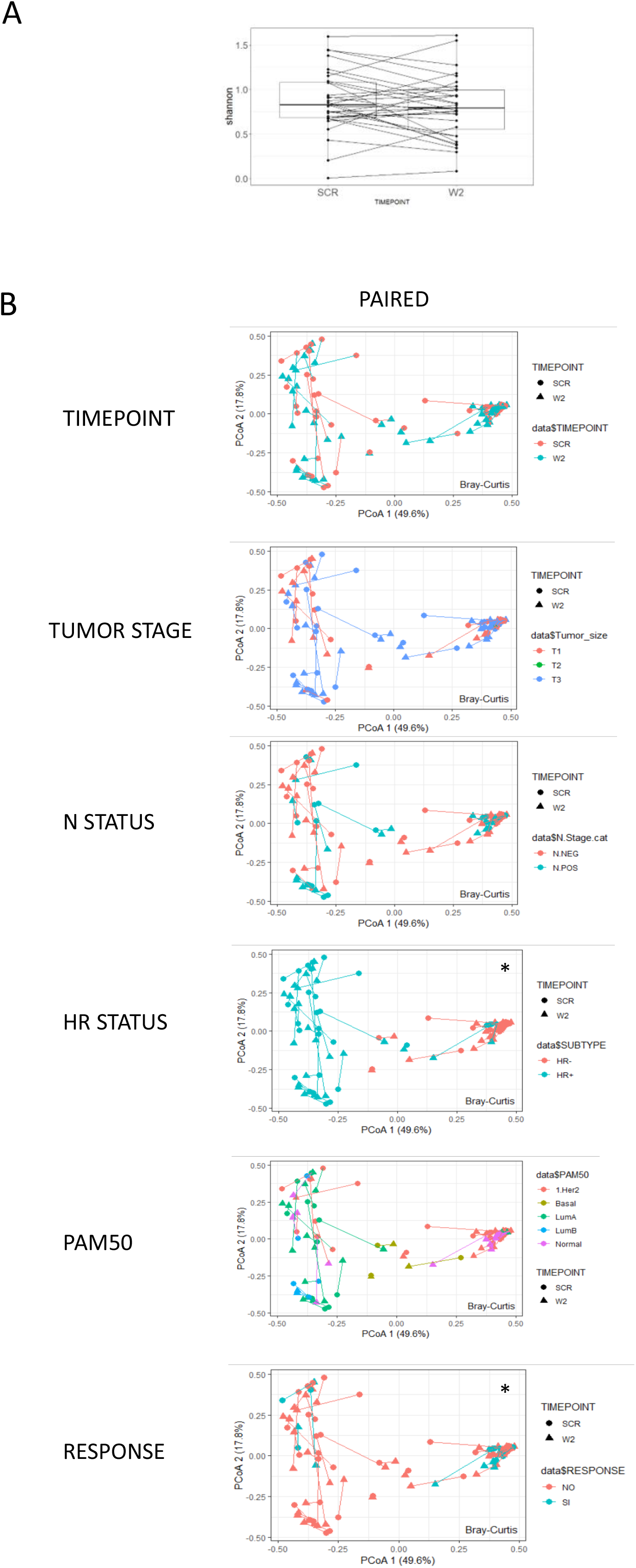
Heterogeneity analyses. **A)** Box plots showing non-significant intratumoral diversity variation in paired samples during the treatment. **B)** PCoA analysis plots of Bray-Curtis computed distances in paired samples. Different clinicopathologic features are shown in different colors. Star shows the groups that are significantly different in phenotype composition.

**Supplementary Figure 3.**
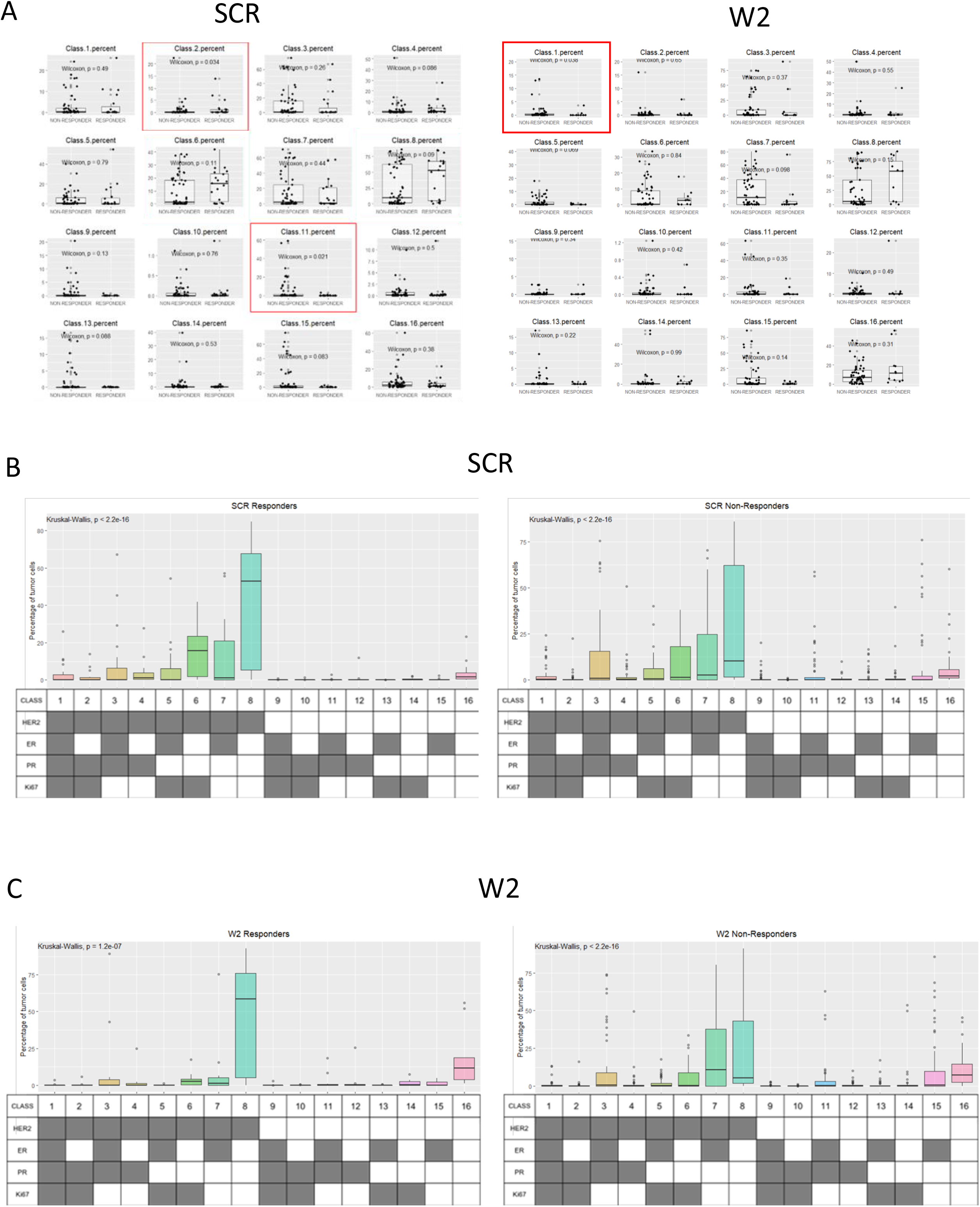
Results from response analyses. **A)** Box plots of the different phenotypes proportions in responder and non-responder patients in SCR and day-15 samples. **B)** Box plots of the different phenotype proportions on responder and non-responder SCR samples. C) Box plots of the different phenotype proportions on responder and non-responder day-15 samples.

**Supplementary Figure 4.**
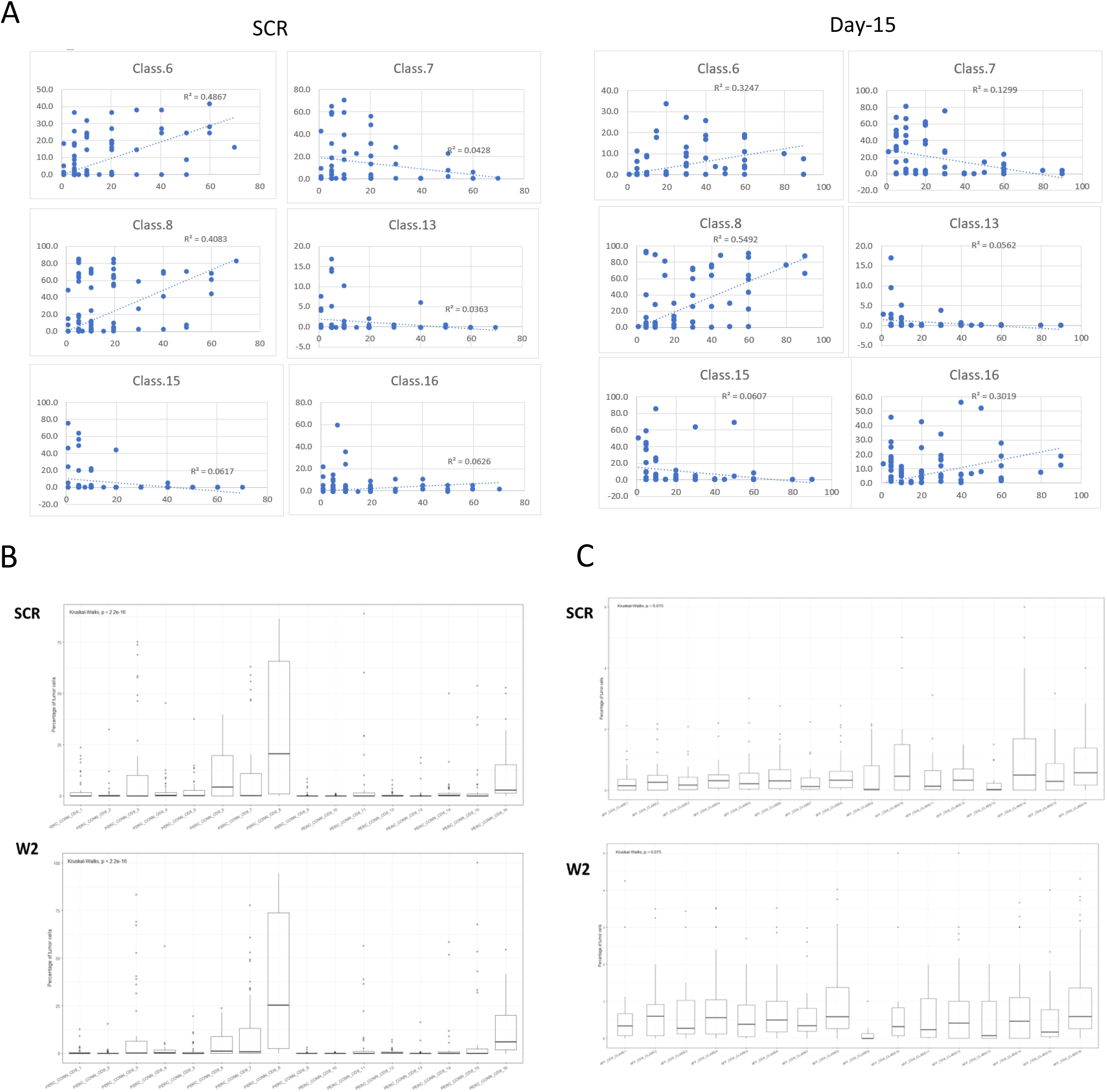
Spatial analyses. **A)** Scatter plot and Spearman correlation coefficient of TILs and different phenotypes in SCR and day-15 simples. B) Box plots of CD8 connections with the different tumor phenotypes for SCR and day-15 samples. C) Box plots of CD8 affinity to the different tumor phenotypes for SCR and day-15 samples.

